# Group size effects in social evolution

**DOI:** 10.1101/319350

**Authors:** Jorge Peña, Georg Nöldeke

## Abstract

How the size of social groups affects the evolution of cooperative behaviors is a classic question in evolutionary biology. Here we investigate group size effects in the evolutionary dynamics of games in which individuals choose whether to cooperate or defect and payoffs do not depend directly on the size of the group. We find that increasing the group size decreases the proportion of cooperators at both stable and unstable rest points of the replicator dynamics. This implies that larger group sizes can have negative effects (by reducing the amount of cooperation at stable polymorphisms) and positive effects (by enlarging the basin of attraction of more cooperative outcomes) on the evolution of cooperation. These two effects can be simultaneously present in games whose evolutionary dynamics feature both stable and unstable rest points, such as public goods games with participation thresholds. Our theory recovers and generalizes previous results and is applicable to a broad variety of social interactions that have been studied in the literature.

## 1 Introduction

Cooperative behaviors increase the fitness of other individuals, possibly at the expense of a personal fitness cost (Sachs et al., 2004). Biological examples include the production of extracellular public goods in microbes (e.g., iron-scavenging molecules, West and Buckling 2003, bacteriocins that eliminate competition, Bucci et al. 2011, and factors that contribute to biofilm formation, Rainey and Rainey 2003), vigilance and sentinel behavior in meerkats (Clutton-Brock et al., 1999), group hunting in social carnivores (Packer and Ruttan, 1988), and the costly punishment of free-riders in humans (Raihani and Bshary, 2011). Identifying the different pathways that allow cooperative behavior to be favored by natural selection (Lehmann and Keller, 2006; Nowak, 2006; West et al., 2007; Van Cleve and Akcay, 2014) is important for understanding the origin of social groups (Krause and Ruxton, 2002) and the major transitions in evolution (Maynard Smith and Szathmary, 1995; Bourke, 2011).

Group size is a crucial variable of social life. Therefore, how an increase or decrease in group size affects individual incentives to cooperate is a recurrent question across the behavioral sciences. In economics and political science, the “group-size paradox” refers to cases where larger groups are less successful than smaller groups in pursuing their common goals because individuals have a greater incentive to shirk when group size is large (Olson, 1965; Esteban and Ray, 2001). In behavioral ecology, one of the most replicated findings is the negative relationship between group size and level of vigilance in social foragers due to increased predator detection and dilution of predator risk (Elgar, 1989; Roberts, 1996; Beauchamp, 2008). Increasing group size has also been shown to reduce voluntary contributions to public goods (Isaac and Walker, 1988) and reciprocity-based cooperation in multi-person interactions (Boyd and Richerson, 1988). More generally, however, whether or not larger groups are less conducive to cooperation might depend on specific assumptions about group interactions. In particular, instances of positive group size effects have also been reported in the empirical literature (Isaac et al., 1994; Yip et al., 2008; Powers and Lehmann, 2017) and are of significant theoretical interest (Dugatkin, 1990; Shen et al., 2014; Powers and Lehmann, 2017; Cheikbossian and Fayat, 2018).

To study how the size of social groups affects the evolution of cooperation we follow the standard approach of modelling social interactions as symmetric games with two strategies (“cooperate” and “defect”) between several players, i.e., as symmetric multiplayer matrix games (Broom et al., 1997; Gokhale and Traulsen, 2014). Payoffs depend on the own strategy and on the number of co-players choosing to cooperate, possibly in a nonlinear way. Strategies are genetically or culturally transmitted, and populations are large enough that the replicator dynamic (Weibull, 1995; Hofbauer and Sigmund, 1998) provides a reasonable model of evolution. Within this framework, the stable rest points of the replicator dynamic correspond to evolutionary endpoints, while the unstable rest points signpost the basins of attraction of such evolutionary attractors. Many social dilemmas for which cooperation can be maintained without repeated interactions or genetic assortment have been theoretically studied using this or related formalisms during the last decades (Taylor and Ward, 1982; Palfrey and Rosenthal, 1984; Diekmann, 1985; Boyd and Richerson, 1988; Motro and Eshel, 1988; Dugatkin, 1990; Dixit and Olson, 2000; Goeree and Holt, 2005; Bach et al., 2006; Hauert et al., 2006; Archetti, 2009; Pacheco et al., 2009; Souza et al., 2009; Archetti and Scheuring, 2011; Chen et al., 2013; Van Cleve and Lehmann, 2013; Sasaki and Uchida, 2014; Chen et al., 2015; Peña et al., 2015; Chen et al., 2017; De Jaegher, 2017; dos Santos and Peña, 2017; Kaznatcheev et al., 2017).

To obtain our results, we make use of the fact that the gain function determining the direction of selection in the replicator dynamic is a polynomial in Bernstein form (Farouki, 2012). The coefficients of this polynomial are given by the gains from switching (Peña et al., 2014), i.e., the differences in payoff a focal player obtains by switching from defection to cooperation as a function of the number of other cooperators in the group. Our analysis makes essential use of the structure of the gain sequence of the game, which collects such gains from switching. We illustrate our results with examples and discuss how previous results in the literature (either proven using alternative arguments or hinted at by numerical analysis) can be recovered using our approach.

Under the conditions that payoffs do not depend directly on the size of the group and that the number of interior rest points of the replicator dynamics do not change as group size increases, we establish that the proportion of cooperators at both stable and unstable interior rest points decreases with group size. This finding, summarized in Proposition 1 in Section 3.1, is our main result. Proposition 1 implies that two kinds of group size effects are possible in the games we analyze. First, a negative group size effect, as the levels of cooperation at stable polymorphisms decrease with increasing group size. Second, a positive group size effect, as the size of the basin of attraction of the stable rest point with the largest level of cooperation increases as well. Proposition 2 identifies general conditions under which the number of rest points is independent of group size.

Sections 3.2 and 3.3 explore the consequences of these general results for two important particular cases subsuming many of the multiplayer matrix games appearing in the literature studying the evolution of cooperation (e.g., Dugatkin 1990; Weesie and Franzen 1998; Bach et al. 2006; Pacheco et al. 2009; Souza et al. 2009; Archetti and Scheuring 2011). Section 3.2 considers games with gain sequences having a single sign change. For such games the replicator dynamics have a unique interior rest point that is decreasing in group size (Proposition 3). If the sign change is from positive to negative, the interior rest point is stable and the group size effect is negative, as the proportion of cooperators at the interior rest point decreases. Conversely, if the sign change is from negative to positive, the interior rest point is unstable and the group size effect is positive, as the basin of attraction of full defection decreases while the basin of attraction of full cooperation increases. In Section 3.3, we focus on games characterized by “bistable coexistence” (Gokhale and Traulsen, 2014; Pena et al., 2015), i.e., a phase portrait where the unstable interior rest point divides the basins of attraction of the stable interior rest point and full defection. Such a phase portrait is typical of many nonlinear social dilemmas, including those featuring participation thresholds or public goods games with sigmoid production functions (Dugatkin, 1990; Bach et al., 2006; Pacheco et al., 2009; Souza et al., 2009; Archetti and Scheuring, 2011; Peña et al., 2014; Archetti, 2018). For these games there is both a negative group size effect (as the proportion of cooperators at the stable interior rest point decreases) and a positive group size effect (as the basin of attraction of full defection decreases). This result is stated in Proposition 4. Alternatively, an increase in group size can lead to a loss of both interior rest points. This makes the group size effect negative as an increase in group size results in full defection being the only attracting point of the replicator dynamics.

Several models in the literature consider a more complicated dependence of payoffs on group size than the one we consider in our main result. For instance, if the total benefit from cooperating has to be shared among group members (as in standard formulations of the linear public goods game, see, e.g., Boyd and Richerson 1988), then the gains from switching themselves depend on group size. This introduces an additional effect, which might either reinforce or countervail the fundamental group size effect investigated in Section 3. We investigate this additional effect in Section 4 and state counterparts of Propositions 3 and 4 as Propositions 5 and 6. Section 5 discusses and concludes.

## 2 Model

### 2.1 Social interactions

Social interactions take place in groups of equal size *n*. Throughout the paperThroughout the paper, *n* is treated as a parameter that satisfies 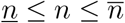 for some given numbers 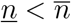 and we use *N* to denote the set of all such group sizes. Individuals within each group participate in a symmetric *n*-player game, playing either strategy *A* (“cooperate”) or strategy *B* (“defect”). The payoff for an individual is determined by its own strategy and the number of other individuals in the group who cooperate but is otherwise independent of group size. Let *a_k_* denote the payoff to an *A*-player (“cooperator”) and *b_k_* denote the payoff to a *B*-player (“defector”) when *k* = 0,1,…, *n* — 1 co-players play *A* (and hence *n* – 1 – *k* co-players play *B*). Irrespective of their own strategy, players prefer other group members to cooperate, i.e., *a*_*k*+1_ ≥ *a_k_* and *b*_*k*+1_ ≥ *b_k_* hold for all *k* = 0,1,…, *n*–2 (Uyenoyama and Feldman, 1980; Kerr et al., 2004). We begin our analysis by assuming that the payoffs *a_k_*, and *b_k_* do not depend explicitly on group size *n*. In Section 4 we relax this assumption.

The gain in payoff an individual makes from cooperating rather than defecting when *k* co-players cooperate is *d_k_* = *a_k_* – *b_k_*. We refer to this as the *k*-th gain from switching (to cooperation). In all of the games we consider in the following, *d_k_* will be negative for some *k*, indicating the presence of a social dilemma in which individuals increase their own payoff by defecting but thereby lower the payoffs of all other group members (Matessi and Karlin, 1984; Kerr et al., 2004).

While our results apply more generally, we will consider a variety of public goods games to motivate and illustrate our results. In these games, cooperators make a costly contribution to the provision of a public good, whereas defectors free ride on the contribution of cooperators. Unless indicated otherwise, we will suppose that the cost *c* > 0 incurred by each contributor is independent of the the number of other contributors and that all group members obtain the same benefit *u_j_*, which is increasing in the number of cooperators *j*. As the number of contributors includes the focal player, we have *j* = *k* if the focal player defects, but *j* = *k* + 1 if the focal player cooperates. Therefore, in such a public goods game payoffs are given by *a_k_* = *u*_*k*+1_ – *c* and *b_k_* = *u_k_*, and the *k*-th gain from switching is *d_k_* = Δ*u_k_* – *c*, where Δ*u_k_* = *u*_*k*+1_ – *u_k_* ≥ 0. Perhaps the simplest example of such a public goods game is the volunteer’s dilemma (Diekmann, 1985) in which at least one cooperator is required for a benefit *v* > *c* to be enjoyed by all group members. This corresponds to the case *θ* = 1 of a threshold public goods game, in which a minimum number *θ* of cooperators is required for a benefit *v* > *c* to be enjoyed by all group members, so that *u_j_* = *v* if *j* ≥ *θ* and *u_j_* = 0 otherwise (Taylor and Ward, 1982; Palfrey and Rosenthal, 1984; Bach et al., 2006; Archetti, 2009).

### 2.2 Evolutionary dynamics

Evolution occurs in a large, well-mixed population with groups of identical size *n* randomly formed by binomial sampling. Hence, if there is a proportion *x* of *A*-players and a proportion 1 – *x* of *B*-players in the population, then the expected payoffs to *A*-players and *B*-players are respectively given by

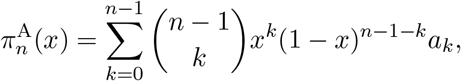

and

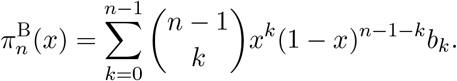

We assume that the change in the proportion of *A*-players over evolutionary time is given by the continuous-time replicator dynamic (Weibull, 1995; Hofbauer and Sigmund, 1998)

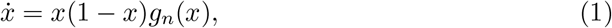

where

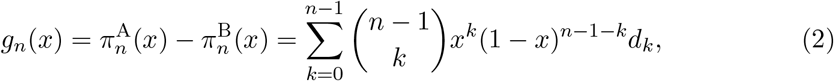

i.e., the difference in expected payoffs between the two strategies, is the “gain function” (Bach et al., 2006), which can also be interpreted as the selection gradient on a continuously varying mixed strategy *x* (Peña et al., 2015). Since the factor *x*(1 – *x*) is always nonnegative, the sign of the gain function *g_n_*(*x*) indicates the sign of *ẋ* in Eq. (1) and hence the direction of selection, that is, whether or not the proportion of *A*-players will increase for a given population composition *x* and group size *n*.

The replicator dynamic has two trivial (or “pure”) rest points at *x* = 0 (where the whole population consists of defectors) and at *x* = 1 (where the whole population consists of cooperators). Interior (or “mixed”) rest points are given by the values *x*\* ∊ (0,1) satisfying

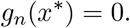

To simplify the exposition, we impose the regularity condition that *_d_g_n___*(*x*\*)/*dx* ≠ 0 holds at all interior rest points. An interior rest point is then stable (i.e., evolutionarily attracting) if and only if *_d_g_n___*(*x*\*)/*dx* < 0 holds, and unstable (i.e., evolutionarily repelling) otherwise. We further suppose that for *n* ∊ *N* the number of sign changes s of the gain sequences (*d*_0_, *d*_1_,…, *d*_*n*–1_) is independent of group size *n* (i.e., the gain sequence 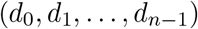 has no sign changes between 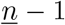 and 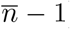), indicating that the fundamental structure of the social dilemma under consideration is the same for all group sizes in the range under consideration. Moreover, we suppose that *s* ≥ 1 holds as otherwise either full defection (*x* = 0) or full cooperation (*x* = 1) is the unique stable rest point of the replicator dynamics for all group sizes *n* ∊ *N*.

## 3 Results

### 3.1 General results

Our first result shows that if the number of interior rest points is independent of group size, then the proportion of cooperators at all interior rest points decreases when group size increases.

#### Proposition 1.

*Suppose that the replicator dynamics* (1)–(2) *have the same number of interior rest points l* ≥ 1 *for all group sizes n* ∊ *N*, *and denote these rest points by* 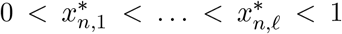 *for group size n. Then* 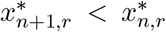 *holds for all* 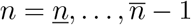 *and r* = 1, …,*l*.

The full proof of Proposition 1 is in Appendix A.1. The key step towards obtaining this result is the following identity, which links the gain functions (and thus the replicator dynamics) for adjacent group sizes:

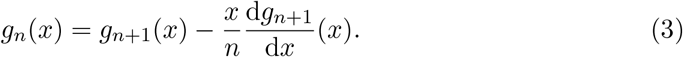

Eq. (3) is a simple consequence of properties of the gain functions *g_n_*(*x*), previously observed by Motro (1991), which stem from the fact that the gain functions are polynomials in Bernstein form (Peña et al., 2014) with coefficients (given by the gains from switching dk) that do not depend on group size.

To see how Eq. (3) yields Proposition 1, observe that this equation implies that at the interior rest points of the dynamic with group size *n* + 1 (where the gain function *g*_*n*+1_(*x*) vanishes), the gain function *g_n_*(*x*) will have the opposite sign of the derivative d*g*_*n*+1_(*x*)/d*x*. This ensures that between any two interior rest points of the replicator dynamic for group size *n* + 1 the replicator dynamic for group size *n* has exactly one rest point. The result then follows upon establishing that the remaining interior rest point for the replicator dynamic with group size n must have a higher proportion of cooperators than the largest interior rest point 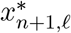 for group size *n* + 1.

**Figure 1:**
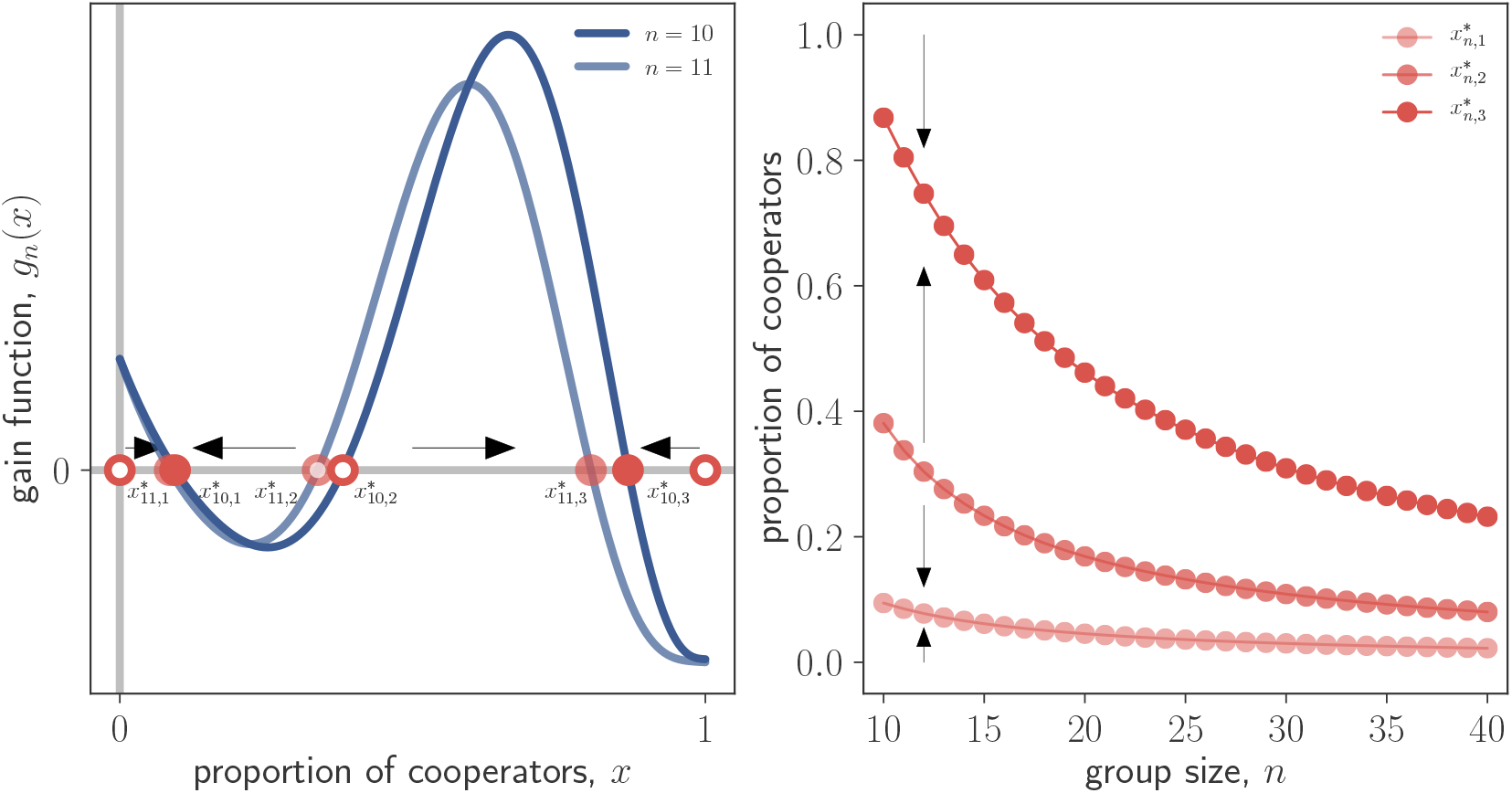
Group size effects in the threshold public good game with an additional reward *δ* > 0 shared among cooperators considered by Chen et al. (2013). Payoffs are given by *a_k_* = *u*_*k*+1_ + *δ*/(*k* + 1) – *c* and *b_k_* = *u_k_*, where *u_j_* = v if *j* ≥ *θ* and *u_j_* = 0 otherwise. In all panels, *c* = 1, *v* = 5, *θ* = 7, *δ* = 1.5. *Left panel:* Gain functions (*blue lines*) with corresponding rest points (*red symbols*), and direction of selection (*black arrows*) for two group sizes: *n* = 10, and *n* = 11. Full circles represent stable rest points and empty circles represent unstable rest points. *Right panel:* Proportion of cooperators at the interior rest points as function of group size for 10 ≤ *n* ≤ 40. The direction of selection (*black arrows*) is also shown. As group size increases, the proportion of cooperators at interior rest points decreases.

The decrease in the proportion of cooperators at all interior rest points as group size increases asserted in Proposition 1 leads to contrasting effects of group size on the evolution of cooperation. First, there is an obvious negative group size effect, as the proportion of cooperators at stable polymorphisms decreases with group size. Second, the proportion of cooperators at unstable rest points decreases as well. As the rest points of the replicator dynamics alternate between being stable and unstable, this implies an increase in the size of the basin of attraction of the stable rest point with the largest proportion of cooperators. Hence, there is also a positive group size effect. These two effects are illustrated in Fig. 1 for the relatively complex case of a game with three interior rest points: 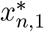 (stable), 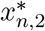 (unstable), and 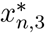 (stable). In line with Proposition 1, larger group sizes lead to smaller proportions of cooperators at the two stable interior rest points 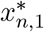 and 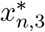 but also, via a decrease in the value of 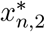, to a larger basin of attraction for 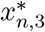 and a smaller basin of attraction for 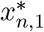. As xn 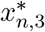 sustains a higher level of cooperation than 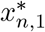, this latter effect can be said to promote the evolution of cooperation.

**Figure 2:**
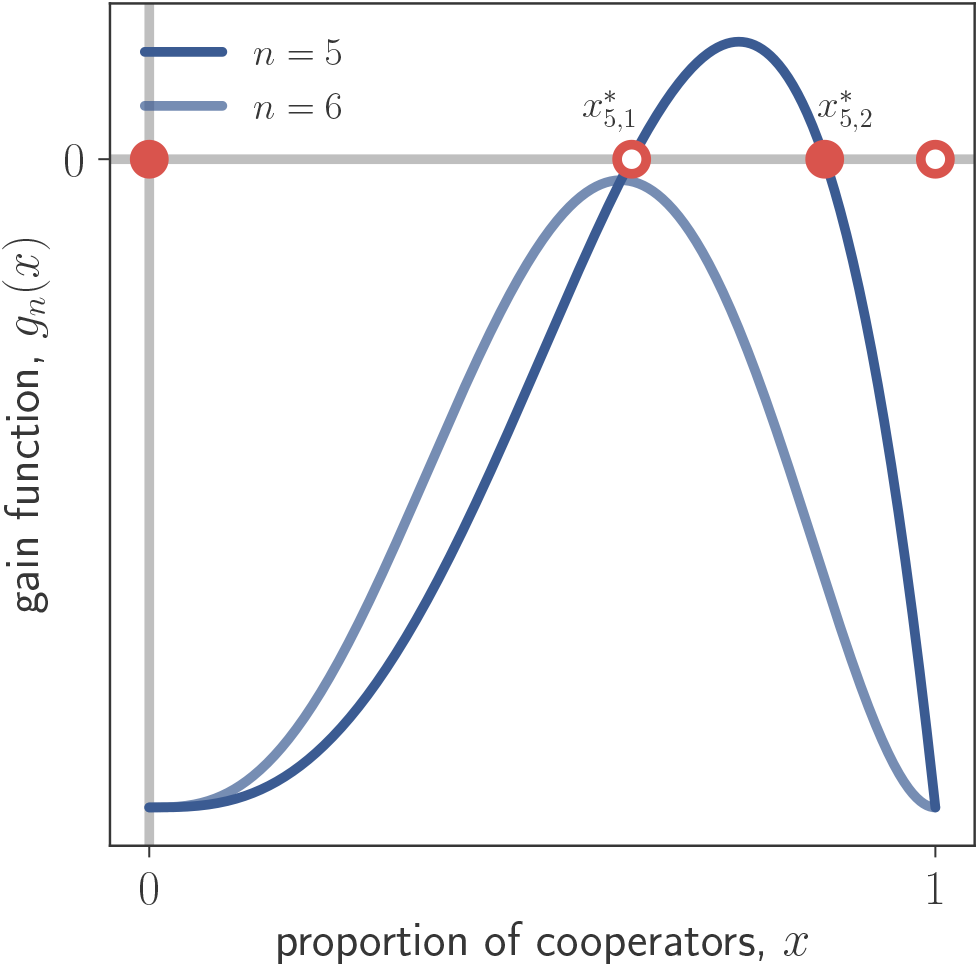
An increase in group size can lead to a reduction in the number of rest points. Here we illustrate this effect for a threshold public goods game with *c* = 1, *v* = 2.8, *θ* = 4, which has two interior rest points for group size *n* = 5 but no interior rest points for group size *n* = 6.

Proposition 1 is predicated on the assumption that the number of interior rest points for the different group sizes under consideration is the same. This does not have to be the case. In particular, it is possible that an increase in group size leads to a decrease in the number of rest points. Fig. 2 illustrates this possibility for the case of a threshold public goods game. On the other hand, the arguments establishing Proposition 1 show that an increase in group size can never lead to an increase in the number of rest points. Further, it is known that the number of interior rest points of the replicator dynamics cannot exceed the number of sign changes s in the gain sequences (Peña et al., 2014, Property 2). Therefore, if the number of interior rest points of the replicator dynamic for the maximal group size 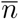 is equal to s, then the number of interior rest points of the replicator dynamics is independent of group size. The proof of the following result in Appendix A.2 demonstrates that, in addition, if an increase in group size causes a reduction in the number of interior rest points, then the number of rest points is reduced by an even amount.

#### Proposition 2.

*Suppose that the number of interior rest points of the replicator dynamic* (1)–(2) *for group size* 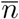 *is equal to the number of sign changes s of the gain sequences. Then for all group sizes n* ∊ *N the number of interior rest points of the replicator dynamics is equal to s. More generally, if* 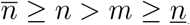, *then the number of interior rest points of the replicator dynamic with group size n is either equal to the number of interior rest points of the replicator dynamic with group size* m *or lower by an even amount.*

### 3.2 Games with a unique interior rest point

Suppose that for all group sizes *n* ∊ *N* the replicator dynamics have a unique interior rest point that, for simplicity, we denote by 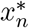. It is then immediate from Proposition 1 that the proportion of cooperators at this rest point is decreasing in group size. Combining this observation with the sufficient condition for the existence of a unique interior rest point from Result 3 in Peña et al. (2014) immediately yields:

#### Proposition 3.

*Suppose that for all n* ∊ *N the gain sequences have a single sign change (s* = 1*). Then the replicator dynamics* (1)–(2) *have a unique interior rest point for all n* ∊ *N, and the proportion of cooperators* 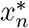 *at this interior rest point is decreasing in group size n*.

Proposition 3 encompasses two cases. First, the gains from switching can be positive for a small number of cooperators (up to some threshold 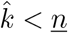) and negative for a large number of cooperators (beyond the threshold 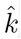). In this case there exists a unique interior rest point 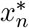 that is also the unique stable rest point of the replicator dynamics (Peña et al., 2014, Result 3.2). For this case, Proposition 3 indicates that the group size effect is negative in the sense that an increase in group size causes a decrease in the proportion of cooperators at equilibrium. This finding generalizes a result due to Motro (1991), who showed that there is a unique stable interior rest point and a negative group size effect for public goods games with concave benefits and intermediate costs (for which Δ*u_k_*, and therefore *d_k_*, is decreasing in *k*, and 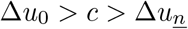 holds). It also generalizes the well-known result that the proportion of cooperators at the unique stable rest point of the volunteer’s dilemma is decreasing in group size (cf., e.g., Archetti 2009) and corresponding results for the volunteer’s dilemma with cost sharing (Dugatkin, 1990; Weesie and Franzen, 1998). This last example, which differs from the other two in that the gains from switching are not monotonically decreasing in k, is illustrated in Fig. 3.

The second case encompassed by Proposition 3 is the one in which the gains from switching are negative for a small number of cooperators (up to some threshold 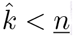) and positive for a large number of cooperators (beyond the threshold 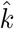). In this case the two trivial rest points *x* = 0 and *x* = 1 are stable and the unique interior rest point 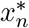, which separates the basins of attraction of the two stable rest points, is unstable (Peña et al., 2014, Result 3.2). For this case, Proposition 3 indicates that the group size effect is positive in the sense that with an increase in group size the basin of attraction of full defection (x = 0) shrinks while the basin of attraction of full cooperation (*x* = 1) increases. For public goods games with convex benefits and intermediate cost (for which Δ*u_k_*, and therefore *d_k_*, is increasing with 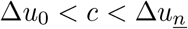) this effect has been previously noted in Motro (1991).

**Figure 3:**
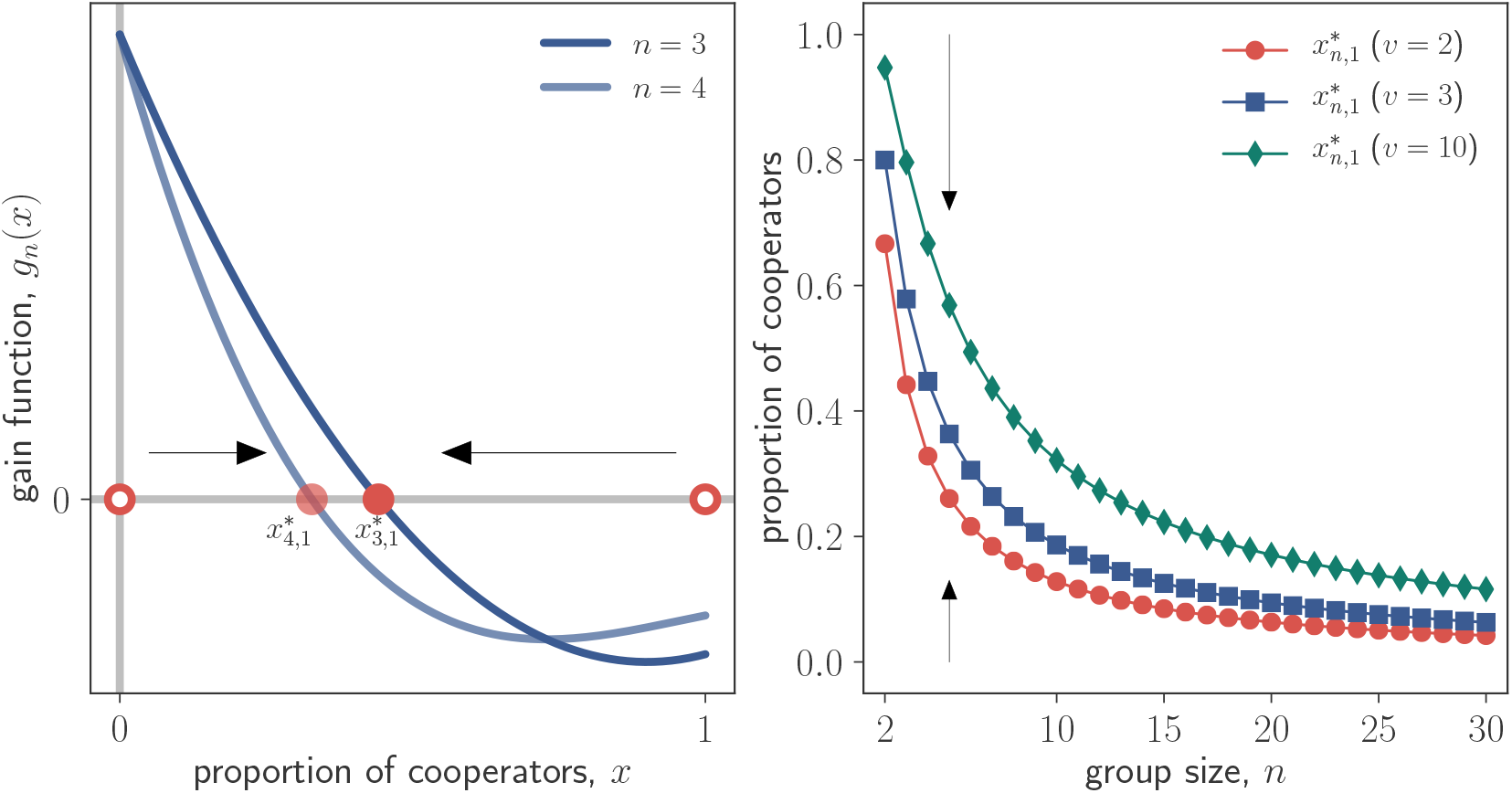
Group size effects in the volunteer’s dilemma with cost sharing considered by Weesie and Franzen (1998). Payoffs are given by *a_k_* = *v* – *c*/(*k* + 1), *b*_0_ = 0, and *b_k_* = *v* for *k* ≥ 1. In all panels, *c* = 1. *Left panel:* Gain functions (*blue lines*) with corresponding rest points (*red symbols*), and direction of selection (*black arrows*) for *v* = 2, and two group sizes: *n* = 3, and *n* = 4. Full circles represent stable rest points and empty circles represent unstable rest points. *Right panel:* Proportion of cooperators at the interior rest point as function of group size for different parameter values. The direction of selection (*black arrows*) is also shown. As group size increases, the proportion of cooperators at the unique stable interior rest point decreases, i.e., the group size effect is negative.

### 3.3 Games with two interior rest points

Many social dilemmas are such that defection is individually advantageous if the number of cooperating co-players is either sufficiently small or sufficiently high, while cooperation is individually advantageous in between, i.e., the gains from switching satisfy *d*_0_ < 0 and the gain sequences (*d*_0_, *d*_1_,…, *d*_*n*–1_) have two sign changes for all group sizes *n* ∊ *N*. This scenario arises in the threshold public goods game when the minimum number θ of cooperators required for the benefit *v* > *c* to be enjoyed by all group members satisfies 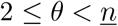. More generally, public goods games in which the benefits from the provision of the public good are sigmoid in the number of contributors and the costs of provision are intermediate (Archetti and Scheuring, 2011; Archetti, 2018) have this structure. Peña et al. (2014) provide further examples and discussion.

Assuming that the gains from switching have the structure described above ensures that the rest point at *x* = 0 is stable and the rest point at *x* = 1 is unstable for all *n* ∊ *N* (Peña et al., 2014, Result 1). Further, there are at most two interior rest points satisfying 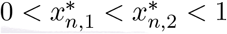, with the smaller of these rest points 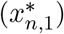 being unstable and the larger interior rest point 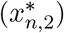 being stable. The existence of these rest points is guaranteed if 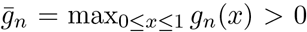 holds (Peña et al., 2014, Result 4.1). Combining these observations with the arguments yielding the results in Section 3.1, Appendix A.3 proves:

#### Proposition 4.

*Suppose that for all n* ∊ *N the gain sequences have two sign changes (s* = 2*), their initial signs are negative, and that* 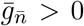 *holds. Then, the replicator dynamics* (1)–(2) *have two interior rest points for all group sizes n* ∊ *N. Further, at both the unstable rest point* 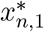 *and the stable rest point* 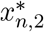 *the proportion of cooperators is decreasing in group size and we have*

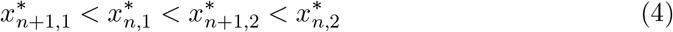

*for all* 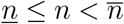.

Proposition 4 indicates that there are two different effects of group size on cooperation in games with two interior rest points. First, there is a negative group size effect, as the proportion of cooperators at the stable interior rest point decreases as group size increases, i.e., 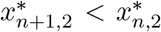 holds. Second, there is a positive group size effect, as the proportion of cooperators at the unstable interior rest point also decreases as group size increases, i.e., 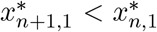 holds, implying that the basin of attraction of full defection (*x* = 0) shrinks while the basin of attraction of the stable rightmost rest point increases. These effects are in line with what happens in games with a unique interior rest point that we have discussed in Section 3.2. The additional twist is that rather than having the group size effect being negative or positive depending on the structure of the game, both the negative and the positive group size effects co-occur in the same game.

Fig. 4 illustrates Proposition 4 for the case of a threshold public goods game. As noted above, the result is applicable more generally. For instance, the observations (obtained via numerical calculations) that both interior rest points decrease with group size for the *n*-person tit-for-tat model of Dugatkin (1990) (his “Model II”) and the *n*-person snowdrift game discussed by Souza et al. (2009), are implied by Proposition 4.

**Figure 4:**
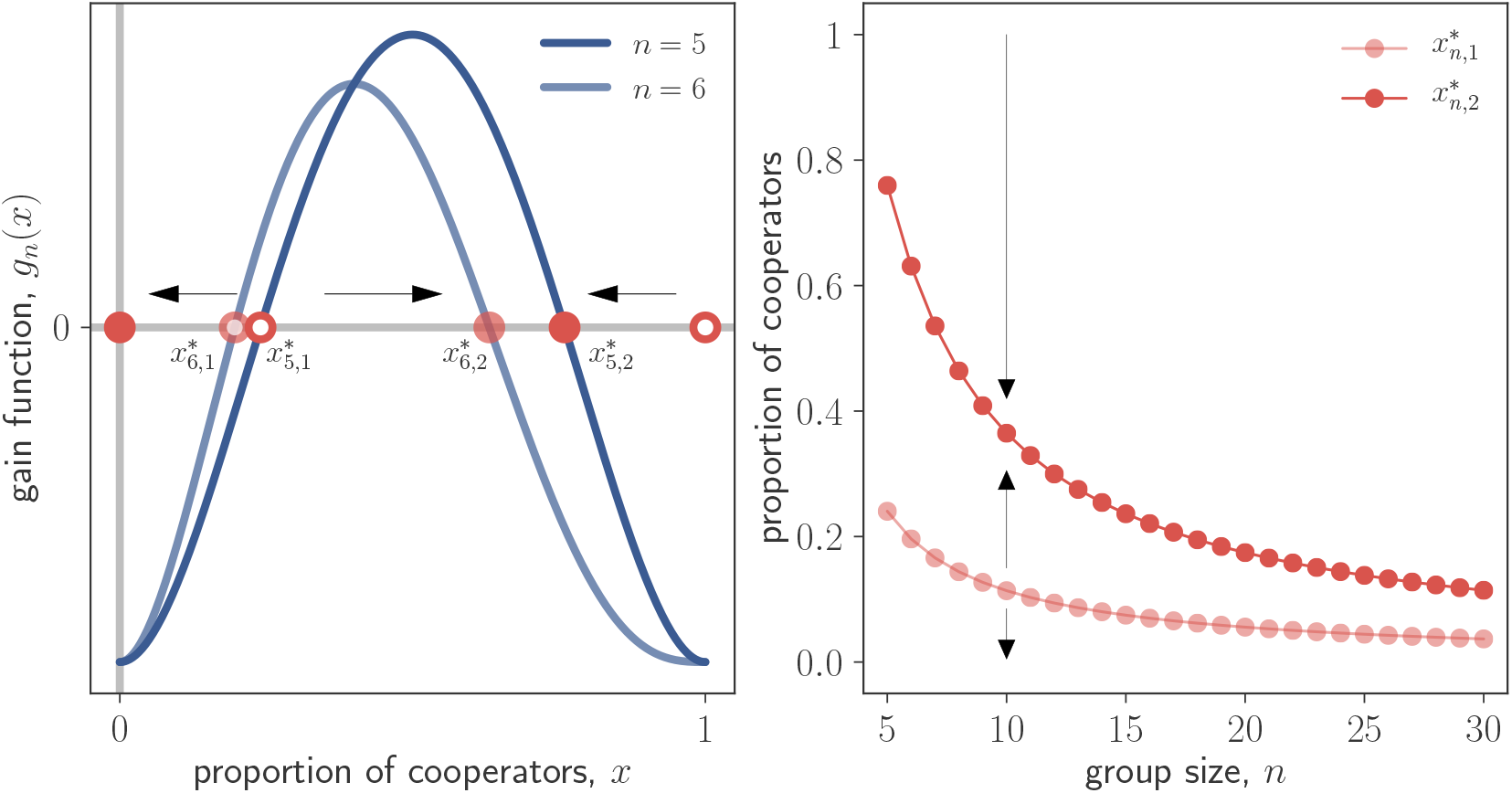
Group size effects in a threshold public good game. Payoffs are given by *a_k_* = *u*_*k*+1_ – *c* and *b_k_* = *u_k_*, where *u_j_* = *v* if *j* ≥ *θ* and *u_j_* = 0 otherwise. In all panels, *c* = 1, *v* = 5, and *θ* = 3. *Left panel:* Gain functions (*blue lines*) with corresponding rest points (*red symbols*), and direction of selection (*black arrows*) for two group sizes: *n* = 5, and *n* = 6. Full circles represent stable rest points and empty circles represent unstable rest points. *Right panel:* Proportion of cooperators at the interior rest points as function of group size for 5 ≤ n ≤ 30. The direction of selection (*black arrows*) is also shown. As group size increases, the proportion of cooperators at both interior rest points decreases. This leads to both a negative group size effect (the proportion of cooperators at the interior stable rest point decreases) and a positive group size effect (the basin of attraction of the interior stable rest point increases).

The role of the condition 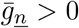 in the statement of Proposition 4 is to ensure that the replicator dynamic has two interior rest points for group size 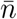 and, therefore, has these two rest points for all group sizes (Proposition 2). If the reverse inequality 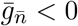 holds, then for large groups there are no interior rest points, whereas (provided that the inequality 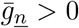 holds) for small group sizes there exists two interior rest points. In such a situation there is (as illustrated in Fig. 2) a critical group size such that for smaller group sizes the rest point *x* = 0 is the only stable rest point, whereas for larger group sizes there is a stable polymorphism at which some proportion of the population cooperates. Hence, this describes a case in which the group size effect is unambiguously negative.

## 4 Extension: games with gain sequences depending on group size

So far our analysis has assumed that the payoffs *a_k_* and *b_k_*, and therefore the gains from switching *d_k_*, depend only on the number of other cooperators a focal player interacts with and not directly on the size of the group. This assumption is not always warranted. For instance, Hauert et al. (2006) and Pacheco et al. (2009) consider variants of a public goods game in which the benefits *u_k_* from cooperation are shared among all group members rather than accruing to each individual. The payoffs to cooperators and defectors are then 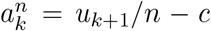 and 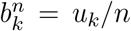. The resulting gains from switching

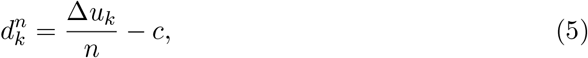

depend not only on *k* but also on group size *n*.

If the gains from switching are, as in Eq. (5), decreasing in group size, then the proportion of cooperators at an unstable interior rest point may increase with group size. In particular, as illustrated for the case of a threshold public goods game in Fig. 6 below, Propositions 3 and 4 are no longer applicable to describe the group size effect on unstable interior rest points. Further, there is no hope to obtain a counterpart to Proposition 1. In the following we therefore focus on stable interior rest points and show that for these the conclusions from Propositions 3 and 4 remain intact.

Consider, first, the case in which the gain sequences 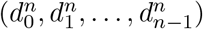 have a single sign change from positive to negative for all group sizes n ∈ N. This ensures that the replicator dynamics, given by Eq. (1) with

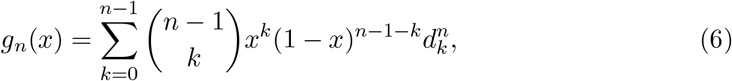

have a unique interior rest point 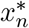 for all group sizes *n* ∊ *N* and that this rest point is stable. Appendix A.4 shows the following result, and Fig. 5 illustrates it for the model with discounted benefits proposed by Hauert et al. (2006).

**Figure 5:**
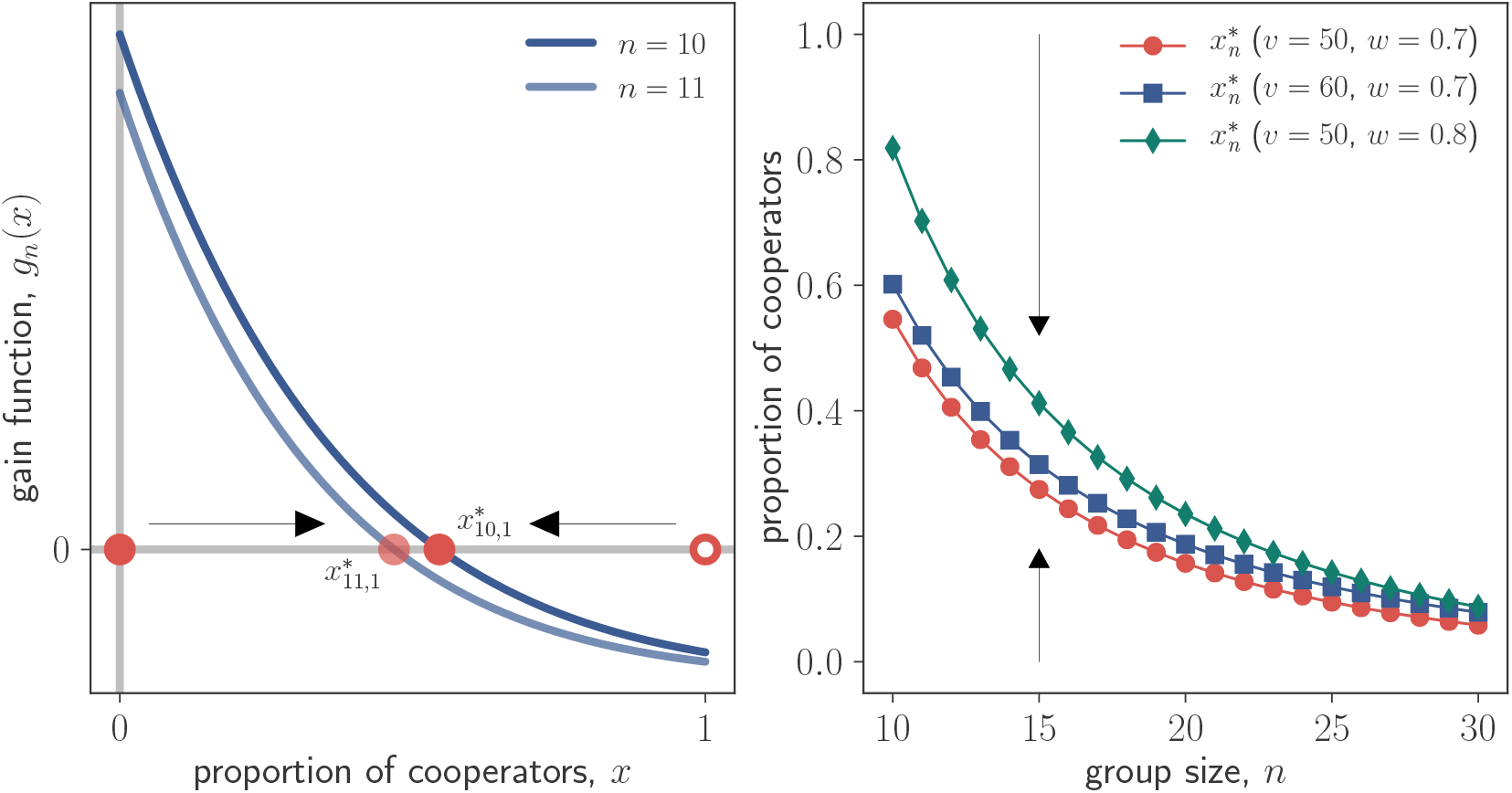
Group size effects in the model with discounted benefits from Hauert et al. (2006). Payoffs are given by *a_k_* = *u*_*k*+1_/*n* – *c* and *b_k_* = *u_k_*/*n*, where *u_k_* = *v*(1 – *w^k^*)/(1 – *w*) with 0 < *w* < 1. For intermediate values of 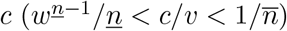 the gains from switching 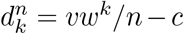 satisfy the assumptions in the statement of Proposition 5. In all panels, *c* = 1. *Left panel:* Gain functions (*blue lines*) with corresponding rest points (*red symbols*), and direction of selection (*black arrows*) for *v* = 50, *w* = 0.7, and two group sizes: *n* = 10, and *n* = 11. Full circles represent stable rest points and empty circles represent unstable rest points. *Right panel:* Proportion of cooperators at the interior rest point as function of group size for different parameter combinations. The direction of selection (*black arrows*) is also shown.

### Proposition 5.

*Suppose that the gain sequences* 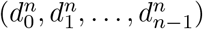 *have a single sign change from positive to negative for all n* ∊ *N. Then the replicator dynamics defined by* *Eq.* (1) *and* *Eq.* (6) *have a unique stable interior rest point* 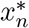 *for all n* ∊ *N. Further, if* 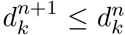 *holds for all k* = 0,1,…, *n* – 1 *and all n satisfying* 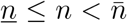, *then the proportion of cooperators* 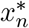 *at this interior rest point is decreasing in group size n.*

The intuition for Proposition 5 is that a decrease in the gains from switching decreases the gain function and that such a decrease in the gain function reduces the proportion of cooperators at the stable interior rest point. Therefore, the negative dependence of the gains from switching on group size considered here reinforces the group size effect observed in Proposition 3 by further reducing the proportion of cooperators at the stable interior rest point. The same intuition applies to the following counterpart to Proposition 4 that we prove in Appendix A.5:

### Proposition 6.

*Suppose that the gain sequences* 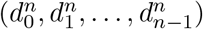 *have two sign changes, their initial signs are negative, and that* 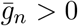 *holds for all n* ∊ *N. Then the replicator dynamics defined by* *Eq.* (1) *and* *Eq.* (6) *have two interior rest points* 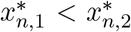 *for all group sizes n* ∊ *N with the first of these unstable and the second stable. Further, if* 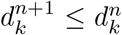 *holds for all k* = 0,1,…, *n* – 1 *and all n satisfying* 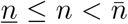, *then the proportion of cooperators at the stable rest point* 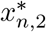 *is decreasing in group size n.*

Fig. 6 illustrates the conclusions from Proposition 6 for a game having the same structure as a threshold public goods game, except that the benefits *u_k_* are shared among all group members as in Eq. 5. Fig. 6 also illustrates that sharing the benefits among more group members decreases the gain function and thereby increases the proportion of cooperators at the unstable interior rest point compared to the benchmark case considered in Proposition 4. Depending on parameter values, this effect may or may not be large enough to overturn the conclusion from Proposition 4.

## 5 Discussion

We have investigated how group size affects the evolutionary dynamics of multiplayer cooperation. More specifically, we have shown that an increase in group size can have a negative effect (a decrease in the proportion of cooperators at equilibrium) and a positive effect (an increase in the basin of attraction of the stable rest point sustaining the largest proportion of cooperators) on social evolution. Depending on the payoff structure of the social interactions one effect can be present and the other absent (as in games featuring a single interior rest point), or both effects can be present at the same time (as in games featuring two interior rest points). For threshold public goods games and other games characterized by bistable coexistence both the invasion barrier needed for cooperators to invade a population of defectors and the proportion of cooperators expected at the stable interior rest point decrease as group size increases. We have also shown that if payoffs depend explicitly on group size and such dependence is negative, the negative group size effect is reinforced, while the positive group size effect is attenuated or, depending on the particular payoff structure of the game, reversed.

**Figure 6:**
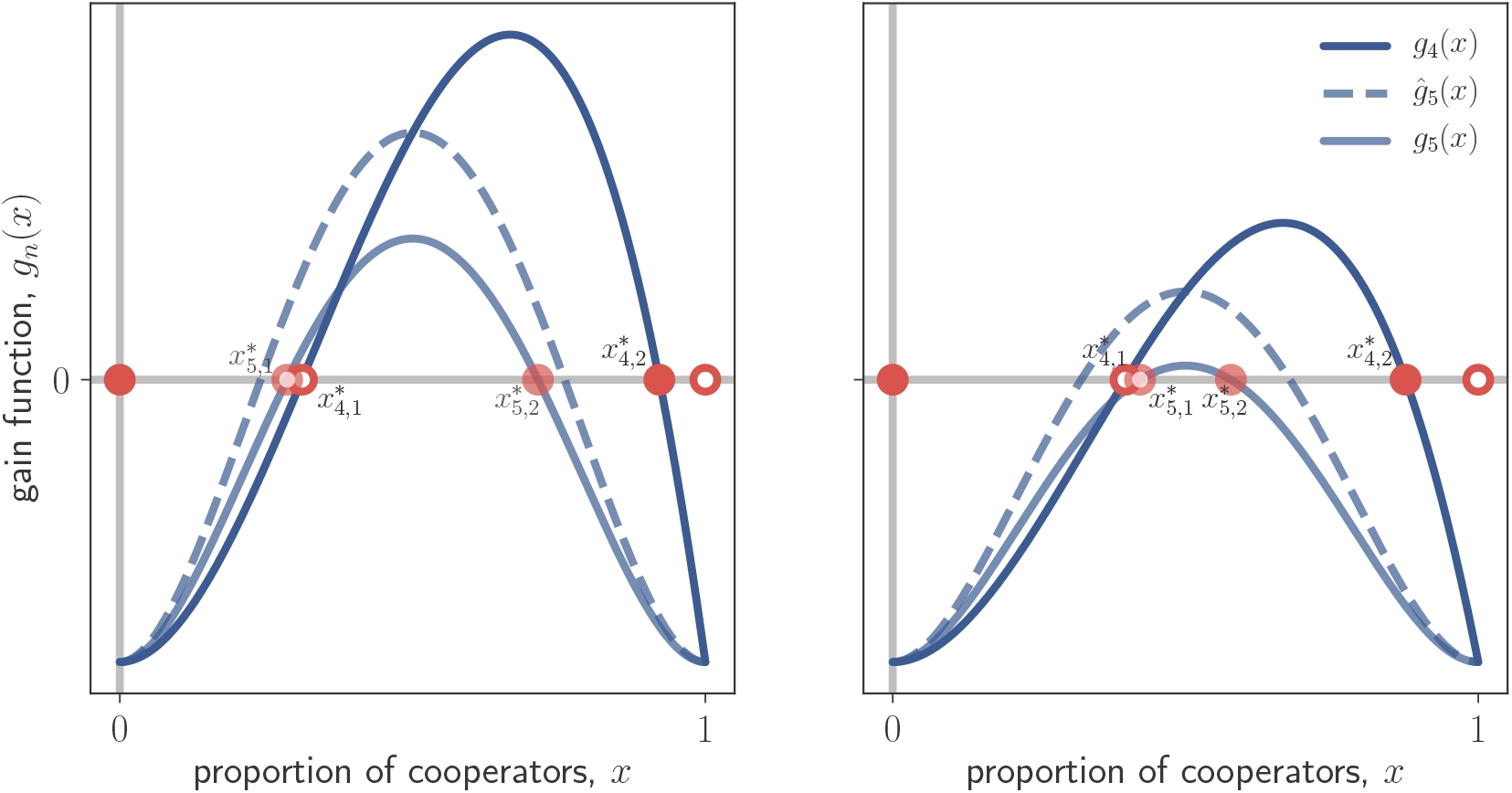
Illustration of Proposition 6 for the case of a threshold game with shared benefits. Payoffs are given by 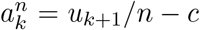 and 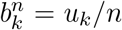, where *u_j_* = *v* if *j* ≥ 3 and *u_j_* = 0 otherwise. Both panels show the gain functions *g_n_*(*x*) for group sizes *n* = 4 and *n* = 5 (*solid lines*) and the gain function 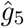*(x)* for the larger group size (*dashed lines*) corresponding to the benchmark of a threshold public goods game in which payoffs for group size 5 are the same as for group size 4. In both panels the proportion of cooperators at the stable interior rest point decreases as group size increases. *Left panel:* the proportion of cooperators at the unstable interior rest point decreases as group size changes from *n* = 4 to *n* = 5 (*v* = 20, *c* = 1). *Right panel:* the proportion of cooperators at the unstable interior rest point increases as group size changes from *n* = 4 to *n* = 5 (*v* = 14, *c* = 1).

The negative group size effect we identify is in line with the common expectation that the selection pressure on certain types of cooperation decreases as group size rises. Such a negative group size effect requires that the gain sequence is sometimes decreasing, meaning that individual incentives to cooperate are (at least for some social contexts) decreasing in the number of cooperators in the group. When this is the case, the decisions to cooperate are strategic substitutes (Bulow et al., 1985); equivalently, cooperation is discounted or subject to diminishing returns. Anti-predator vigilance often follows this payoff structure, as the presence of other vigilant individuals usually disincentivizes individual investment in vigilance, i.e., there is a “many eyes” effect (Pulliam, 1973; McNamara and Houston, 1992); in extreme cases one vigilant individual is enough for the group to be protected (Bednekoff, 1997; Clutton-Brock et al., 1999). In agreement with our results, empirical and theoretical studies indicate that vigilant behavior often decreases with group size (Elgar, 1989; McNamara and Houston, 1992; Beauchamp, 2008).

Contrastingly, the positive group size effect we identify has been less emphasized in evolutionary game theory (but see Sumpter and Brännström 2008 and Cornforth et al. 2012, who demonstrate this effect in models with continuous strategies). In an early paper, Dugatkin (1990) noted that, in his model of n-person reciprocity, the threshold frequency of cooperators needed to invade a population of defectors decreased as group size increased. Our analysis reveals that such a positive group size effect is not specific to the payoff structure assumed in Dugatkin (1990), but that it holds more generally for any matrix game featuring unstable interior rest points. As a necessary condition for the existence of unstable interior rest points is that the gain sequence is sometimes increasing, the group size effect can be positive only when the individual incentives to cooperate are (for at least some social contexts) increasing in the number of cooperators in the group. In this case, the decisions to cooperate are strategic complements (Bulow et al., 1985); equivalently, cooperation is synergistic or subject to increasing returns. A common form of synergistic cooperation occurs when a critical number of cooperators is required for cooperation to be individually worthwhile. Examples of such threshold effects have been documented in empirical studies, and hypothesized to be a causal factor behind inverse density dependence or Allee effects (Courchamp et al., 1999). For instance, a large critical number of bark beetles is needed to overcome the defenses of the tree they attack (Franceschi et al., 2005), and cooperative hunting often requires a critical number of hunters to be energetically efficient (Creel and Creel, 1995; Alvard and Nolin, 2002; MacNulty et al., 2014). Also, in group-hunting sailfish, a larger number of hunters improves the hunting success of the group by allowing individuals to alternate their attacks (Herbert-Read et al., 2016), and by keeping group-level unpredictability high in the face of individual lateralization (Kurvers et al., 2017). In all of these cases of synergistic cooperation, our theory suggests that larger groups can be more favorable to cooperation and less favorable to free riding. Indeed, this general prediction is in agreement with both general models of synergistic cooperation with continuous cooperative investments (Cornforth et al., 2012), and a recent mechanistic model of free riding in group-hunting sailfish (Herbert-Read et al., 2016).

We used a variety of public goods games to illustrate our results. In such games, both cooperators and defectors gain equal access to the collective good produced by cooperators, i.e., the collective good is public. Notwithstanding the importance of these models, there are other social dilemmas for which public goods games are not a natural description of the relevant strategic trade-offs. For instance, social interactions can take the form of a collective action problem where the produced good can be accessed only by cooperators or only by defectors, i.e., the collective good is in some sense excludable. Group size effects in such “club” and “charity” goods games (Peña et al., 2015) are readily amenable to analysis by applying our results.

We conclude by noting that our analysis assumed populations were well-mixed and hence without genetic structure. This assumption is not always justified, as many social interactions take place in spatially structured populations characterized by non-negligible amounts of genetic structure (Rousset, 2004; Lehmann and Rousset, 2010; Van Cleve, 2015). A simple way of modeling social evolution in these populations is to focus on a continuously varying mixed strategy and to identify the convergence stable strategies of the resulting adaptive dynamics (e.g., Rousset 2004; Van Cleve and Lehmann 2013; Peña et al. 2015). In this case, the counterpart to the gain function we have analyzed in this paper is also a polynomial in Bernstein form, now with coefficients given by “inclusive gains from switching” depending on the payoffs of the game, the group size, and demographic parameters of the particular spatial model determining the degree of genetic relatedness and the amount of local competition (Peña et al., 2015). In this light, the analysis conducted here is also relevant to investigate group size effects in genetically structured populations, provided that the likely dependence of the inclusive gains from switching on group size is taken into account. Investigating the effects of group size on the evolution of cooperative behaviors under nontrivial population structure with the tools developed here would complement recent efforts in this area (Shen et al., 2014; Powers and Lehmann, 2017; Van Cleve, 2017).

## Acknowledgements

J. Peña gratefully acknowledges financial support from the ANR-Labex IAST.

## Appendix

## A.1 Proof of Proposition 1

We first obtain Eq. (3). To do so, we make use of two identities established in the appendix of Motro (1991). Using our notation for the gain function and the gains from switching, these identities are

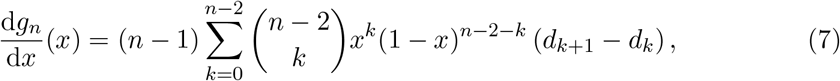

and

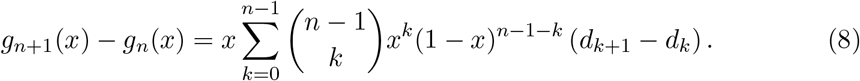

Applying Eq. (7) (which is nothing but the derivative property of polynomials in Bernstein form; see, e.g., Peña et al. 2014) to group size *n* + 1 and dividing both sides of the resulting equation by n yields

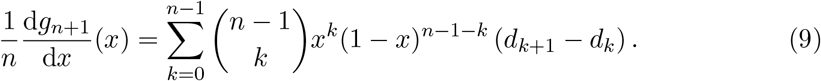

Substituting Eq. (9) into Eq. (8) we obtain

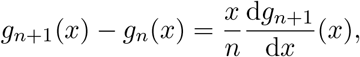

from which Eq. (3) is immediate.

Consider any *n* satisfying 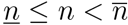. The following establishes that the replicator dynamic for group size *n* must have a rest point in the interval 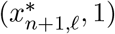: Because *g*_*n*+1_(*x*) has no root in 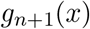, *g*_*n*+1_(*x*) has the same sign as the derivative 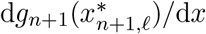 for all 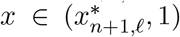. As the gain sequences (*d*_o_,…, *d*_*n*–1_) and (*d*_o_,…, *d_n_*) have the same initial sign (given by the sign of the first non-zero gain from switching *d_k_*) and the same number of sign changes *s*, they also have the same final sign. Hence, the final sign of the gain sequence (*d*_0_,…, *d_n_*) is the same as the sign of 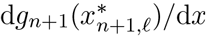, too (Peña et al., 2014, Property 1). Therefore, for sufficiently large 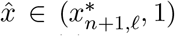 the sign of 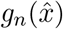 coincides with the sign of 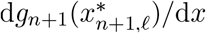. From Eq. (3) and 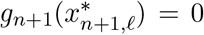 we then have that 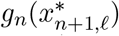 and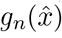 have opposite signs, so that *g_n_*(*x*) has a root in the interval 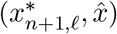. Consequently, the replicator dynamic for group size *n* has a rest point in the interval 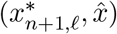. For the case *l* = 1 this finishes the proof of the proposition.

Suppose *l* ≥ 2 and let *n* again satisfy 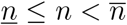. Consider (with *r* = 1,…,*l* – 1) any adjacent interior rest points 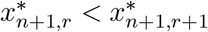 of the replicator dynamic for group size *n*+1. As stable and unstable rest points alternate, the derivatives 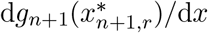 and 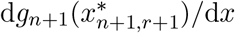 have opposite signs. As 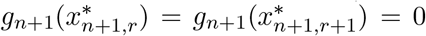 holds, it follows from Eq. (3) that 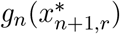 and g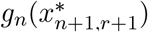 have opposite signs, too. Therefore, *g_n_*(*x*) has at least one root in the interval 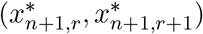, with each such root corresponding to an interior rest point of the replicator dynamic for group size *n*. As there are *l* – 1 intervals of the form 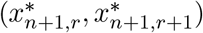 and the replicator dynamic for group size *n* has an interior rest point in the interval 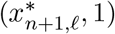, this implies that there is exactly one interior rest point of the replicator dynamic for group size *n* in each of the intervals 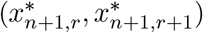 for *r* = 1,…, *l* – 1. Therefore, for all 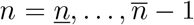 and *r* = 1, …, *l* – 1, we have

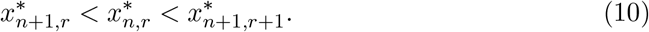

In conjunction with the inequality 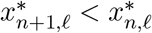 established in the preceding paragraph, Eq. (10) finishes the proof.

## A.2 Proof of Proposition 2

Let *l* denote the number of interior rest points of the replicator dynamic for a given group size *n* ∊ *N*. We begin by showing that the number of interior rest points of the replicator dynamic for group size *m* ∊ *N* satisfying *m* < *n* must be at least *l*. This is trivially true for *l* = 0, so consider *l* ≥ 1. By the same arguments as in the proof of Proposition 1, the replicator dynamic for group size *n* – 1 has at least one rest point in the interval 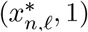 and, in case *l* > 1, at least one rest point in each of the intervals 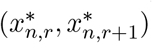 for *r* = 1,…,*l* – 1. As there are *l* – 1 such intervals, the replicator dynamic for group size *n* – 1 has at least as many rest points as the replicator dynamic for group size n. By a straightforward induction argument, it follows that the same conclusion obtains not only for group size *n*– 1 but for all group sizes *m* ∊ *N* satisfying m < n.

Suppose that the number of interior rest points for group size n is equal to the number of sign changes s of the gain sequences. It then follows from the argument in the previous paragraph that, for all group sizes *n* ∊ *N*, the number of interior rest points is at least s. On the other hand, the number of interior rest points of the replicator dynamic for group size n cannot be larger than the number of sign changes s of the gain sequence (Peña et al., 2014, Property 2). Hence, independently of group size the number of interior rest points is s.

The assumption that the regularity condition d*_g_n__*(*x*\*)/d*x* ≠ 0 holds for all interior rest points implies that all roots of the polynomials g_n_(x) are simple. Therefore, for all group sizes *n* ∊ *N* the number of interior rest points is either equal to the number of sign changes s of the gain sequences or less by an even amount Peña et al. (2014, Property 2). It follows that the number of interior rest points for the replicator dynamics for two different group sizes either are equal or differ by an even amount. As it has been established above that the number of interior rest points cannot increase with group size, this observation finishes the proof.

## A.3 Proof of Proposition 4

From Result 4.1 in Peña et al. (2014) the condition 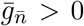 (in conjunction with the assumption on the sign pattern of the gain sequences) is sufficient to imply that the replicator dynamic for group size 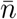 has two interior rest points 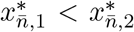 with the first of these being unstable and the second stable. As the gain sequences have two sign changes for all *n* ∊ *N*, Proposition 2 then implies that the replicator dynamic for any group size *n* ∊ *N* has two interior rest points with the same stability pattern. From Proposition 1, the inequalities 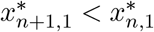 and 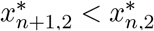 hold for all *n* satisfying 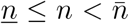. The remaining inequality in Eq. (4) follows from Eq. (10) in the proof of Proposition 1 in Appendix A.1.

## A.4 Proof of Proposition 5

The existence of a unique interior rest point 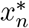 and its stability for all group sizes is immediate from Result 3 in Peña et al. (2014).

Fix *n* satisfying 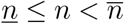 and let

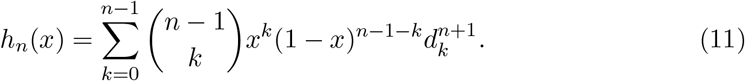

Observe that the assumption 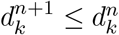 for all *k* = 0,1,…, *n* – 1 implies *h_n_*(*x*) ≤ *g_n_*(*x*) for all *x* ∊ [0,1], where *g_n_*(*x*) has been defined in (6).

An argument identical to the one that we have used to obtain Eq. (3) in Appendix A.1, yields

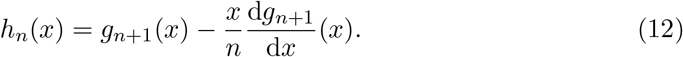

As the rest point 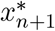 is stable, Eq. (12) implies 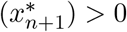 and therefore 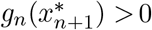. By the stability of the rest point 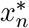, we have *g_n_*(*x*) < 0 for all 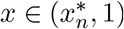. Therefore, the inequality 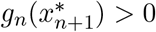 implies 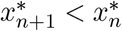, which is the desired result.

## A.5 Proof of Proposition 6

From Result 4.1 in Peña et al. (2014) the condition 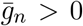 (in conjunction with the assumption on the sign pattern of the gain sequences) is sufficient to imply that for all group sizes *n* ∊ *N*, two interior rest points 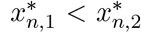 exist with 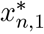 being unstable and 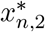 being stable. The proof is then finished by observing that the same argument as in the proof of Proposition A.4 implies the inequality 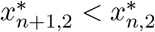 for all *n* satisfying 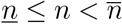.

